# Compatible Comma-Free Codes and the Origin of the Genetic Code

**DOI:** 10.64898/2026.10.08.757769

**Authors:** Latham Boyle, Lillian Boyle

## Abstract

The genetic code is the rule that (apart from minor modifications [1, 2]) is used by all contemporary life on Earth to translate genes into proteins. It is summarized by a 4 *×* 4 *×* 4 table which maps the 64 possible triplet RNA codons to the 20 canonical amino acids (“AAs”). Understanding the origin of this code is a fundamental open problem in biology. We review the idea of a comma-free code, and put forward a general reason to suspect the primordial code was comma-free (even though the modern code is not). We explain why this primordial code should then persist as a comma-free subcode within the modern code, and identify the five optimal candidates for such a subcode. We then introduce the idea of “*compatible* comma-free codes” and explain how they allow multiple codes to evolve in parallel, circumventing a standard argument suggesting that code evolution should effectively freeze when the associated genome becomes sufficiently complex. In particular, we highlight the possibility that the original code was a comma-free doublet code, which was then overtaken by a compatible comma-free triplet code, which finally evolved (after the evolution of the start codon and ribosome) to the modern saturated triplet code. To assess this idea, we classify all inequivalent compatible comma-free doublet-triplet codes and, among these, identify the three most promising predecessors of the modern code. Consistency between these three best codes and the “wobble rules” in gene translation provide non-trivial evidence for this possibility.

## I. INTRODUCTION

The genetic code is the rule that translates genes (expressed as strings of the four RNA bases U,C,A,G) into proteins (expressed as strings of the 20 canonical acids (“AAs”)). This rule, which is used – apart from minor modifications [1, 2] – by all current life on Earth, may be summarized in a famous 4 *×* 4 *×* 4 table (Fig. 2). Understanding the origin of this code is a fundamental open problem in biology. The code’s universality implies it dates to the early history of life on Earth (predating our last universal common ancestor [3, 4]), and progress on understanding its origins might be hoped to yield new clues about the beginnings of life itself.

Many clues and theories have been put forward about the origin of the code: see [5–74] and references therein for an introduction to the large literature on this topic. It should be emphasized that these various suggested theories are, in many cases, not mutually exclusive, but may rather be different pieces of a common puzzle; and the same goes for the new observations we offer below.

First, in Section II, we start by reviewing what it means for a code to be *comma-free* [75–77] (Fig. 1) and explaining why this is a natural property to expect of the *primordial* code (even though it is not true of the *modern* code). We explain how a primordial comma-free code can evolve into the modern *saturated* code and why, in this case, the primordial comma-free code should still be embedded as a *subcode* of the modern genetic code. We find that the modern code contains five optimal subcodes (Fig. 2). These form our initial list of best candidates for the comma-free predecessor of the modern code.

**FIG. 1.**
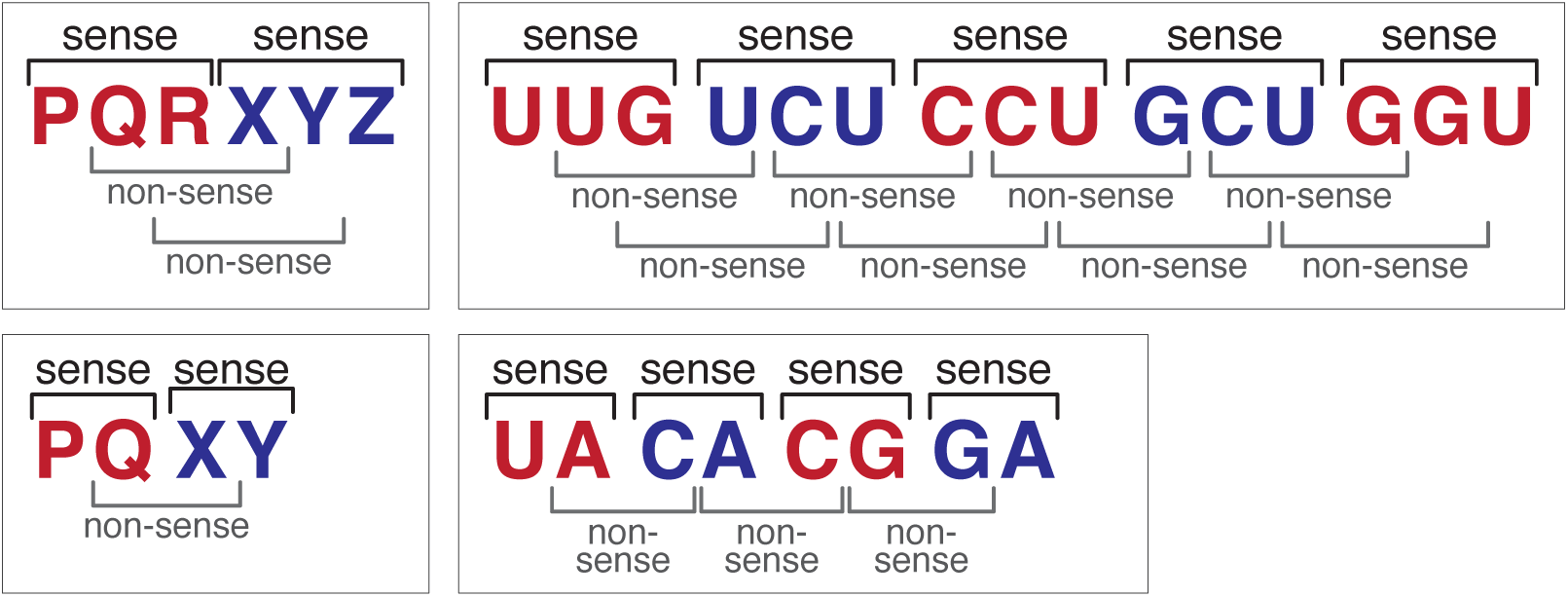
Comma-free conditions for a triplet code (top row) and a doublet code (bottom row).

**FIG. 2.**
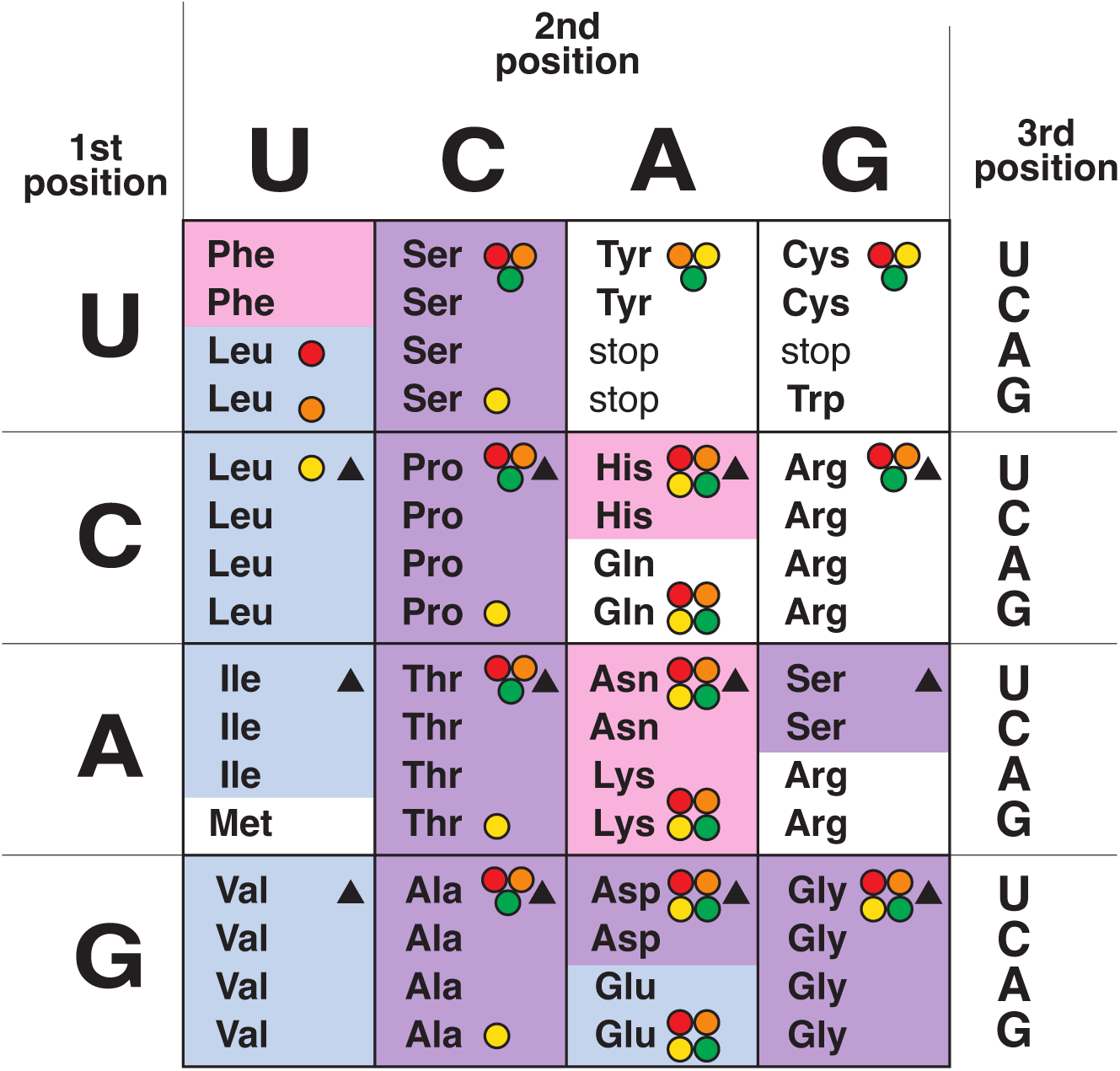
The 5 optimal comma-free subcodes within the modern genetic code. As in all figures that follow, the blue region shows the AAs that occur prebiotically, the pink region shows the AAs that correspond to Class-2 aminoacyl tRNA-synthetases, and the purple region shows the six AAs in both categories. The black triangles show the unique comma-free subcode that maximizes the number of prebiotic AAs (9) and then total AAs (12); it also includes 8 Class-2 AAs. The four colors of circles display the four comma-free subcodes that maximize the number of Class-2 AAs (9) and then total AAs (14); three of these codes contain 8 prebiotic AAs and one contains 7.

Next, in Section III we introduce the idea of *compatible* comma-free codes (Figs. 4, 3), and explain how they permit more general evolution of the code, in a way that circumvents a standard argument [78] (associated with “frozen accident hypothesis” [9]) suggesting that the evolution of the code should (nearly) freeze once the corresponding genome becomes sufficiently complex. We explain why, in fact, this freezing argument only applies to a (nearly) saturated code.

Next we explain why evolution from a comma-free doublet code to a compatible comma-free triplet code is of particular interest. Let us call such a pair of compatible codes a “doublet-triplet code” for short. In this scenario, the resulting triplet code should again be embedded as a subcode of the modern code. We distinguish between two possible evolutionary paths, which we call *two-step* and *three-step* doublet-triplet evolution; and, in each case, we then enumerate the optimal comma-free doublet-triplet predecessors of the modern code. In particular, the three optimal three-step doublet-triplet codes (which turn out to be a subset of the five optimal subcodes found in Sec. II) form our refined list of best candidates for the comma-free predecessor of the modern code.

### Finally, in Section IV, we discuss several key points about the results of the previous sections

First, we explain how to extend the notion of comma-free code to incorporate the phenomenon of “wobble” in gene translation, and point out that nearly every sufficiently long code that would be comma-free if the standard Watson-Crick pairing rules were strictly obeyed, fails to remain comma-free once wobble effects are taken into account. We then emphasize how striking it is that the three best codes found in Sec. III (and, more generally, most of the various optimal codes found in Secs. II and III) turn out to *remain* comma-free even when wobble effects are taken into account.

We explain why this provides non-trivial evidence for the possibility that our modern code had a comma-free triplet predecessor (and, perhaps even for the possibility that this, in turn, had a compatible comma-free doublet predecessor).

We end by considering the possibility of other, even simpler starting points (which turn out to not be promising from the standpoint of our compatible-comma-free line of reasoning), and mentioning some topics for future research.

## II. COMMA-FREE CODES

In this section, we first introduce the idea of comma-free codes and the reason that it is natural to expect the primordial code to be comma-free. We then show that there are five optimal comma-free subcodes embedded within the current code.

### A. Why comma-free codes?

Soon after the discovery of DNA in 1953 [79, 80], it was realized by Gamow and others [81] that the genetic code operated by splitting a string of DNA bases into triplet “codons,” like a string of 3-letter words with the spaces removed. This led to a question. Since a given string of letters may be grouped into triplets in three different ways:

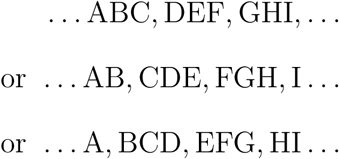

how does the cell know which of these three “reading frames” is the correct one when reading a gene?

In 1957, Ref. [75] proposed an answer: the genetic code might be *comma-free*. A comma-free code is one in which only a subset of the 4^3^ = 64 possible triplets are *valid* (or *“sense codons”*), while the remaining triplets are *invalid* (or *“nonsense codons”*); and, moreover, two adjacent sense codons never combine to yield another intermediate (overlapping) sense codon (see Fig. 1). These conditions ensure that the correct reading frame may be inferred, without the need for any additional punctuation (“commas”) to indicate how the letters should be split into triplets. In more concrete terms, if we imagine that each RNA sense codon^1^ has a corresponding tRNA molecule whose anticodon binds to it, while each nonsense codon does not, these conditions ensure that the tRNA anticodons will only bind to the RNA in the “right” frame, and never bind in either of the two “wrong” frames. Ref. [75] also showed that, in a four-letter comma-free triplet code, at most 20 of the 64 possible triplets can be sense codons.

When the modern genetic code was decoded experimentally in the 1960s, we learned it is *not* comma-free. On the contrary, it is a *saturated code* in the sense that nearly all of the 64 possible triplets are sense codons (with only three nonsense codons: the stop codons). Still, as emphasized by Crick [9], although the modern code is not comma-free, the *primordial* code may have been.

Indeed there is a basic reason to suspect the primordial code was comma-free rather than saturated. The central puzzle about the origin of life is how some system could have transitioned (with sufficient probability) from inorganic, thermodynamic disorder, to the minimal amount of organized complexity needed to sustain reproduction and evolution. In this context, the fundamental reason that a comma-free code is more plausible than a saturated code is that, in a comma-free code the correct reading frame is specified *locally* by the gene itself, whereas in saturated code it must be specified *remotely*, and then *remembered* locally by some extrinsic coordinating mechanism. This is illustrated by the modern code, where the correct reading frame is determined remotely by a start codon at one end of a gene, and then the ribosome must accurately remember this frame as it walks all the way from one end of the gene to the other (a distance of *∼* 10^3^ bases, say). This mechanism is clearly too sophisticated to be primordial, but in a saturated code, some such sophistication is a logical necessity, since the gene does not contain the relevant information locally. By contrast, in a comma free code, the problem is solved automatically, since a (proto-)tRNA molecule can *only* bind to the gene in the correct frame!

### B. Comma-free code evolution

In this subsection, let us understand three key points about the expected evolution of the genetic code table itself:

1. If a genome is sufficiently complex, the corresponding genetic code is expected to only (or dominantly) evolve by converting nonsense codons to sense codons.
2. Thus, the primordial comma-free code would be expected to initially have evolved by expanding its set of sense codons while *maintaining* the comma-free condition; and then, after the invention of (proto-versions of) the start codon and ribosome, it would have continued to evolve by expand its set of sense codons *without* maintaining the comma-free condition, therefore relaxing to a saturated code (*i.e.* the modern code).
3. Thus, the maximal primordial comma-free code (as existed just before the invention of start codon and ribosome), is expected to persist as a maximal comma-free subcode embedded within the modern saturated code.

Let us now unpack these three points:

**P1oint 1.** Let us start by understanding why, when a genome is sufficiently complex, the corresponding genetic code only evolves by expanding its repertoire of sense codons (*i.e.* by converting nonsense codons to sense codons).

When we say the code maps a certain sense codon *c* to a certain amino acid *x*, we mean that there is a tRNA (or proto-tRNA) adaptor molecule, with “one end” that binds to *c*, and the “other end” that incorporates amino acid *x* in the corresponding protein. Let us schematically denote such an adaptor by (*c, x*), and let *N_c_*denote the total number of times that the codon *c* appears in the genome (or, more correctly, in the coding part of the genome). With this notation, let us systematically consider the four code evolution routes, which we call 1a, 1b, 2a, 2b, and analyze in turn:

- i) *Route 1a:* (*c, x*) *→* (*c, x^′^*) *replacement*. In this route, a mutation converts *x → x^′^* at the adaptor’s amino acid end, yielding a new adaptor (*c, x^′^*) that *replaces* the previous adaptor (*c, x*). Thus, each time codon *c* appears in the genome, the new adaptor translates it as *x^′^* rather than *x*, effectively producing *N_c_* simultaneous mutations. For a sufficiently complex genome (with *N_c_ ≫* 1), such a mutation will be lethal, so code evolution will not proceed via this route.
- ii) *Route 1b:* (*c, x*) *→* (*c, x^′^*) *augmentation*. In this route, a mutation converts *x → x^′^* at the adaptor’s amino acid end, yielding a new adaptor (*c, x^′^*) *in addition to* the previous adaptor (*c, x*). Now each time codon *c* appears in the genome, the two competing adaptors translate it in two competing ways – one as *x* and one as *x^′^*. This again effectively produces *N_c_* simultaneous mutations, but instead of each such mutation occurring with probability one it occurs with probability *p*: the probability that the given occurrence of *c* in the genome is translated by (*c, x^′^*) rather than (*c, x*). But the result is the same: assuming the genome is sufficiently complex (*N_c_ ≫* 1 sufficiently large), such a mutation will be lethal, and code evolution will not proceed via this route.
- iii) *Route 2a:* (*c, x*) *→* (*c^′^, x*) *replacement*. In this route, a mutation converts *c → c^′^* at the adaptor’s anti-codon end, yielding a new adaptor (*c^′^, x*) that *replaces* the previous adaptor (*c, x*). Thus, each time codon *c* appears in the geneome, it is now untranslated (*i.e.* it has been converted from sense to nonsense), effectively producing *N_c_* simultaneous mutations (at least^2^) so, for *N_c_ ≫* 1, such a mutation will be lethal, and code evolution will not proceed via this route.
- iv) *Route 2b:* (*c, x*) *→* (*c^′^, x*) *augmentation*. In this route, a mutation converts *c → c^′^* at the adaptor’s anti-codon end, yielding the new adaptor (*c^′^, x*) *in addition to* the previous adaptor (*c, x*). Thus, each time *c* appears in the genome, the adaptor (*c, x*) translates it as before; but – if *c^′^* was already a sense codon with associated adaptor (*c^′^, x^′^*) – then each time *c^′^* appears in the genome, the new adaptor (*c^′^, x*) competes with the previous adaptor (*c^′^, x^′^*) to translate it either as *x* or *x^′^*. (Without loss of generality, we can assume *x* and *x^′^* are different since, otherwise, this mutation would have no effect on the code table.) Thus, if *c^′^ was* already a sense codon, this effectively produces *N_c_′* simultaneous mutations so, for *N_c_′ ≫* 1, such a mutation will be lethal, and code evolution will not proceed via this route. However, if *c^′^* had been a non-sense codon to compete with, then there was no prior adaptor (*c^′^, x^′^*) to compete with, and hence no obstruction to evolution in this route.

In summary, once a genome is sufficiently complex, it is only expected to evolve by adding new sense codons to its previous repertoire of sense codons (via Route 2b).

For this reason, comma-free codes (which contain many available nonsense codons) can still undergo considerable evolution, while fully saturated codes like the modern code (with essential no nonsense codons, apart from a small number of designated stop codons that play an important and widespread role in gene transcription, and are therefore no longer available for conversion to sense codons) are effectively frozen (though minor, slower changes can still occur under special circumstances [1, 2, 88, 89]).

**Point 2.** How then could the code evolve from a primordial comma-free code to the modern saturated code? From the nature of the transition, and using the previous point, we can infer that the evolution had the following stages:

- *Stage 1: comma-free expansion.* In the initial stage, the code relied on its comma-free character to determine the correct reading frame. Therefore, it could only incorporate new sense codons if they continued to satisfy the comma-free constraint.
- *Stage 2: invention of start codon and ribosome.* At some point, the organism must have developed the modern mechanism of determining the reading frame – *i.e.* it must have invented the start codon (or a proto-version) to establish the reading frame, and the ribosome (or proto-version) to remember this frame as it walks from one end of a gene to the other. A natural guess is that this new scheme evolved because it made the translation process more efficient and error-free, but the details of how, when, and why this transition took place are not needed for our argument. The only salient point is that, after this new mechanism developed, the comma-free constraint was no longer necessary to maintain orderly gene translation.
- *Stage 3: further expansion toward saturation.* In this stage, the code can now expand its repertoire of sense codons, without regard to the comma-free restriction, so it expands until it is saturated (or nearly saturated, since at least one codon must still be reserved to play the role of stop codon).

**Point 3.** Crucially, we see from the previous point that the final comma-free code at the end of Stage 1 (just before the invention of the start codon, and the switch to the modern method of determining the reading frame) should still be preserved as a comma-free *subcode* embedded in the modern genetic code.

Moreover, it is expected to be a *maximal* comma-free subcode (meaning that it is not contained within any larger comma-free subcode) since, after the end of Stage 1, the unrestricted expansion of the code in Stage 3 is likely to quickly conflict with the comma-free restriction.

### C. The 5 optimal comma-free subcodes

We have seen that, if the modern code evolved from a comma-free code, that earlier code is expected to be embedded in the modern code as a comma-free subcode. To identify the most promising candidate subcodes, we make use of the following two clues:

1. The 20 canonical AAs may be divided into 10 that are readily produced abiotically (*e.g.* they are found in meteorites, and are readily produced in Miller-Urey-type experiments) [26, 55, 61–65, 90–97], and 10 that are not (they are only produced by living organisms). Let us call these these classes prebiotic and postbiotic. Presumably the earliest AAs incorporated in the code were a subset of the prebiotic class (not necessarily the whole class), whereas the members of the postbiotic class evolved and were incorporated into the code later.
2. Each of the 20 AAs has a corresponding aminoacyl tRNA-synthetase (or “aaRS”): this is the enzyme that binds the AA to its corresponding tRNA molecule. In 1990, it was discovered that the aaRSs are split into two distinct structural classes (Class 1 and Class 2) [98, 99]; and hence their corresponding AAs are split accordingly: 10 in Class 1 (Val, Leu, Ile, Met, Glu, Gln, Trp, Tyr, Arg, and Cys) and 10 in Class 2 (Gly, Pro, Ala, Thr, Ser, His, Asp, Asn, Lys and Phe). For a number of reasons, the Class 2 synthetases are believed to be more ancient [35, 48, 50, 100] and so, again, a natural guess is that the earliest AAs incorporated in the code were some subset of Class 2, whereas the members of Class 1 were incorporated into the code later.

This suggests that the six AAs that are *both* prebiotic *and* in Class-2 (namely: Ala, Asp, Gly, Ser, Pro, and Thr) were likely incorporated into the code relatively early. Indeed, this conclusion has often been reached in the literature, by a variety of lines of argument (see *e.g.* [26, 35, 54–56, 100]).

Therefore, in this section we assume that these six AAs (Ala, Asp, Gly, Ser, Pro, and Thr) were incorporated into the code relatively early in its evolution – in particular, some time during its comma-free phase – *i.e.* during “Stage 1,” before the introduction of the start codon as the mechanism for determining the reading frame. (Note that it is also consistent with this assumption that, by various lines of analysis in the literature, the modern start codon “Met” entered the code relatively late [26, 55].) Within the paradigm that we are investigating in this section (that the code began with a comma-free phase, before entering a later start-codon-based phase), this seems to be a conservative assumption (and it leads us to candidate subcodes that will have other evidence in their favor, as we will discuss later); but we emphasize that it would be valuable for future work to repeat the analysis without this assumption.

We have then carried out an exhaustive search for the optimal comma-free subcodes, with the following results, shown in Fig. 2:

1. A comma-free subcode can encode at most 14 distinct AAs. There are four subcodes achieving this, shown by the four colored circles in Fig. 2. These four codes are then also automatically optimal in the sense that they encode the highest possible number of distinct Class-2 AAs (namely, 9 of 10). (They also encode many of the prebiotic AAs: 8 of 10 for red, orange, and yellow, or 7 of 10 for green.)
2. There is one other notable subcode. Note that a comma-free subcode can encode at most 9 of the 10 prebiotic AAs; and such a subcode can encode at most 12 distinct AAs in total. The unique code which achieves this is shown by the black triangles in Fig. 2. (It also encodes 8 of the 10 Class-2 AAs.)

These five codes form our initial list of the most promising candidates for the primordial comma-free subcode. In the next section, additional considerations will allow us to narrow the list further.

## III. COMPATIBLE COMMA-FREE CODES

In this section, we introduce the idea of *compatible* comma-free codes and explain how they allow an organism to have two (or more) genetic codes, which can evolve in parallel, with one overtaking the other. In particular, we show how this mechanism allows an initial primordial comma-free *doublet* code to be overtaken by a subsequent comma-free triplet code (which can then further expand into the modern saturated code, after the invention of the start codon, as explained in the previous section). We distinguish two paths by which the doublet code could have evolved to the modern code (we call these the two-step or three-step path, respectively) and, in each case, identify the optimal candidates for comma-free predecessors of the modern code; and we draw particular attention to the three optimal three-step predecessor codes.

### A. Characterizing compatible codes

In the previous section, we explained why, for a *single* comma-free genetic code (with sufficiently complex genome), the code table is only expected to evolve by adding sense codons (*i.e.* converting some of the 64 triplets from nonsense to sense).

In this section, though, we point out a more general possibility: a single organism may simultaneously employ two (or more) *different* genetic codes (Code 1, Code 2, …, Code *n*) which interact in parallel with two (or more) distinct non-overlapping subsets its genetic material (Region 1, Region 2, …, Region *n*).

In order for this situation to have been relevant to primordial code evolution, we need a *local* mechanism to ensure the adaptor molecules of Code *i* (which are meant to interact with the sense codons and genes in Region *i*) do not instead interact with the codons and genes in Region *j*, or vice versa (see Fig. 3), which would wreak havoc in the gene translation process. This is neatly solved by generalizing the notion of a single comma-free code to the concept of *compatible* comma-free codes.

**FIG. 3.**
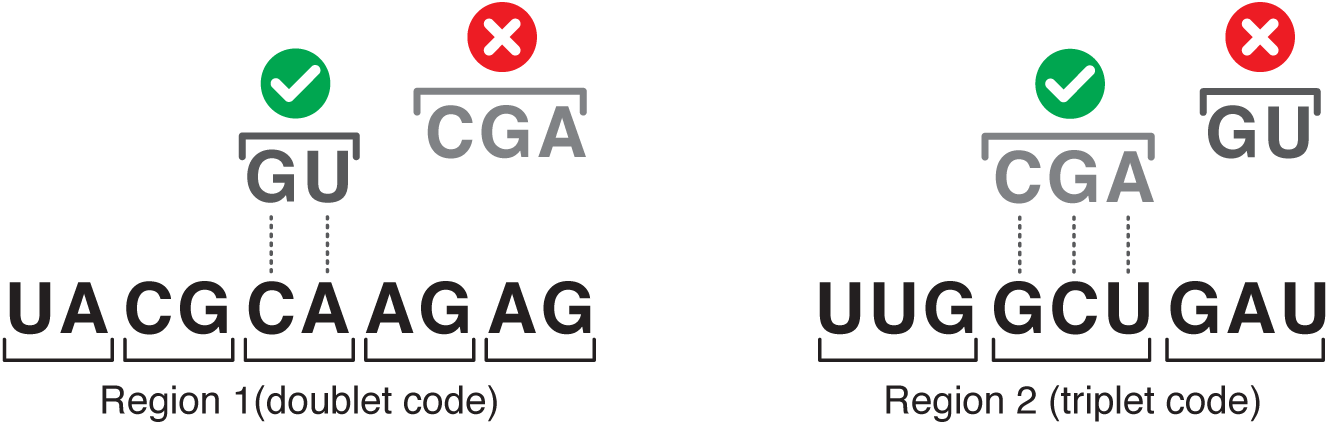
Two hypothetical regions in the primordial genome: Region 1 (comma-free doublet code) and Region 2 (comma-free triplet code). For these codes to not interfere with each other, the sense doublets of Region 1 should never occur in Region 2, and the sense triplets of Region 2 should never occur in Region 1.

**FIG. 4.**
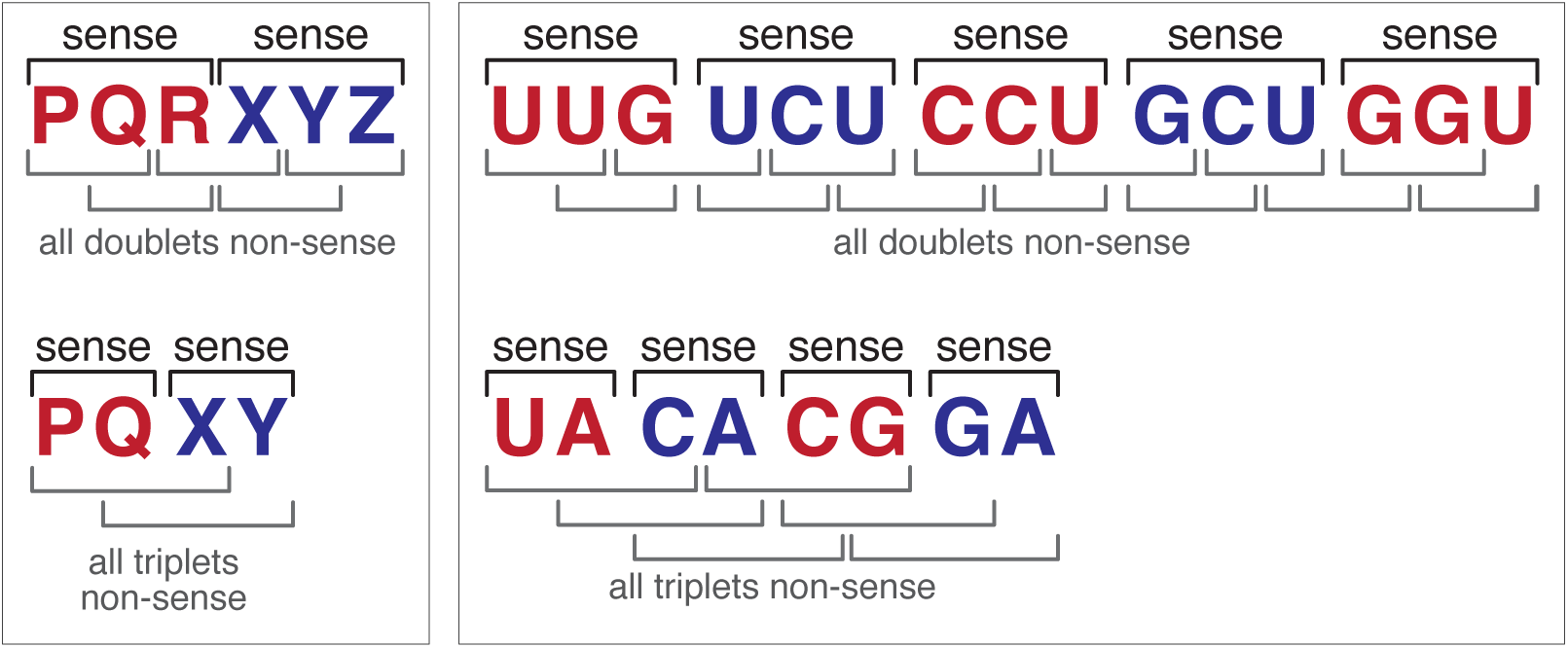
Mutual compatibility conditions between a doublet and triplet code.

We say that two codes (Code *i* and Code *j*) which interact with two regions of the genome (Region *i* and Region *j*) are *compatible comma-free codes* if the following two conditions are satisfied:

1. Code *i* and Code *j* are each comma-free, in the sense that two adjacent sense codons of Code *i* in Region *i* (respectively, of Code *j* in Region *j*) never yield a wrong-frame overlapping sense codon of Code *i* (respectively of Code *j*);
2. Code *i* and Code *j* are mutually compatible in the sense that two adjacent sense codons of Code *i* in Region *i* (respectively, of Code *j* in Region *j*) never produce a sense codon of Code *j* (respectively of Code *i*).

These conditions are illustrated in Figs. 1 and 4 for two mutually-compatible comma-free codes, where one code is a doublet code, and the other is a triplet code. As mentioned in Sec. I, we will refer to such a pair as a “doublet-triplet code” for short.

Let us now consider in more detail how a doublet-triplet code could allow the evolution from a primordial comma-free doublet code to the modern saturated triplet code.

### B. Doublet-to-triplet evolution

Since the early days of the thinking about the code, authors considered the possibility that the current triplet code, which can encode up to 64 amino acids, might have been preceded by a doublet code, which can encode up to 16. Naively, this seems like a natural guess, since at early times there were presumably fewer amino acids to encode, and a doublet code seems like a smaller, simpler starting point.

However, in an early paper on the origin of the genetic code [9], Crick makes a famous argument which seems to rule out the possibility that our current triplet code was preceded by an earlier doublet code. To quote him: “This seems highly unlikely, since it violates the Principle of Continuity. A change in codon size necessarily makes nonsense of *all* previous messages and would almost certainly be lethal.” We now point out a crucial loophole in this argument.

Crick’s conclusion seems unavoidable if we imagine that the old doublet tRNAs and the new triplet tRNAs interact with the *same* stretch of genetic material; or, in other words, if we imagine that the doublet code first learns to make a certain protein, and then we must somehow *convert* the gene for that protein, through a series of random mutations, to the new triplet code (as if we were converting a piece of software to run on a new operating system).

But let us imagine, instead, that the doublet adaptor molecules only interact with a certain subset of the genetic material (“Region 1”), so that the doublet code first learns to make certain proteins on that region; and then, later, the new triplet adaptor molecules (*i.e.* tRNA or proto-tRNA molecules) only interact with a different, non-overlapping subset of the genetic material (“Region 2”), so that the triplet code gradually learns to make new, completely *different* proteins on that region (see Fig. 3).

In this case, the new triplet code could evolve and learn to make increasingly sophisticated proteins on a region that was formerly an unused “junk” region from the standpoint of the doublet code, without interfering with the production of earlier proteins that the doublet code had already learned to make (and could continue making, uninterrupted, as the triplet code evolves). One can imagine that the triplet code might eventually evolve to be superior to the earlier doublet code (both because triplet codons bond more strongly and reliably than doublets, and because triplets can encode a wider variety of AAs), so that the original doublet code becomes vestigial, or disappears altogether.

A key feature of this scenario is that the two codes do not interfere with one another. As explained above, if the two codes form a mutually-compatible pair of comma-free codes (a “doublet-triplet code” for short), this provides an elegant *local* mechanism to ensure this non-interference, without requiring the evolution of any primordially-implausible sophisticated non-local external coordination mechanism.

Now, let us distinguish two different paths by which the evolution from an initial comma-free doublet code to the modern saturated triplet code could have taken place:

- Two-step evolution. In this case:

1. first, a compatible comma-free triplet code evolved to incorporate as many sense codons as it could, while remaining compatible with a given doublet code; and
2. then, with the invention of the modern reading-frame mechanism (start codon and ribosome), this comma-free triplet seed code was freed of its comma-free constraint, and expanded into the modern saturated triplet code (as explained in Sec. II B).
- Three-step evolution. In this case, the evolution contains an extra intermediate step:

1. first, as before, a compatible comma-free triplet code evolved to incorporate as many sense codons as it could, while remaining compatible with a given doublet code;
2. then, the doublet code became vestigial (or disappeared) *before* the evolution of the modern reading-frame mechanism, so that the previous comma-free triplet “seed code” was free to grow further (still constrained to be comma-free, but no longer constrained by compatibility with the doublet code); and
3. finally, with the invention of the modern reading-frame mechanism (start codon and ribosome), this *expanded* comma-free triplet code was freed of its comma-free constraint, and finally expanded into the modern saturated triplet code.

We wish to find the most promising candidates for (both two-step and three-step) doublet-triplet predecessors of the modern code.

### C. Classifying all doublet-triplet codes

As a first step toward identifying the most promising doublet-triplet precessors of the modern code, we begin by classifying all inequivalent compatible doublet-triplet codes (*i.e.* all mutually-compatible pairs of comma-free doublet and triplet codes). We use the generic letters A,B,C,D to denote (some permutation of) the four RNA bases U,C,A,G; and regard two codes as equivalent if they are related by an overall permutation (relabeling) of the letters *{*A,B,C,D*}* or by an overall reflection (*i.e.* a simultaneous reversal of the letters in every sense codon, *e.g.* CAB*→*BAC, ADC*→*CDA).

Given a particular comma-free doublet code *C*_2_ with *n*_2_ sense doublets, a compatible comma-free triplet code *C*_3_ can have at most *n*^max^ sense triplets; we say that if *C*_3_ has this many sense triplets it is *maximal*; and we denote by *N*_ineq_ the number of inequivalent maximal triplet codes *C*_3_ corresponding to the given doublet code *C*_2_.

We have enumerated all inequivalent compatible comma-free doublet-triplet codes, and show the results in Table I, with the notation described above.

**TABLE I.** Inequivalent compatible comma-free doublet-triplet codes. (Notation explained in Sec. III C.)

| $n_2$ | comma-free doublet code | $n_3^{\max}$ | $N_{\text{ineq}}$ | example maximal compatible comma-free triplet code |
| --- | --- | --- | --- | --- |
| 5 | $\{AB, AC, AD, BC, BD\}$ | 2 | 5 | $\{CCD, CDD\}$ |
| | $\{AB, AC, AD, BC, DC\}$ | 2 | 3 | $\{BBD, BDD\}$ |
| 4 | $\{AB, AC, BC, BD\}$ | 6 | 14 | $\{AAD, ADD, BAD, CAD, CCD, CDD\}$ |
| | $\{AB, AC, BD, CD\}$ | 6 | 1 | $\{AAD, ADB, ADD, CAD, CBB, CCB\}$ |
| | $\{AB, AC, AD, BC\}$ | 5 | 14 | $\{BBD, BDD, CBD, CCD, CDD\}$ |
| | $\{AB, AC, DB, DC\}$ | 2 | 5 | $\{AAD, ADD\}$ |
| 3 | $\{AB, AC, DA\}$ | 10 | 28 | $\{ADB, ADC, BBC, BDB, BDC, CBC, CDB, CDC, DDB, DDC\}$ |
| | $\{AB, AC, BC\}$ | 9 | 7 | $\{AAD, ADD, BAD, BBD, BDD, CAD, CBD, CCD, CDD\}$ |
| | $\{AB, AC, BD\}$ | 9 | 7 | $\{AAD, ADB, ADD, CAD, CBB, CCB, CCD, CDB, CDD\}$ |
| | $\{AB, AC, AD\}$ | 8 | 10 | $\{BBC, BBD, CBC, CBD, CCD, DBC, DBD, DCD\}$ |
| | $\{AB, AC, DB\}$ | 8 | 4 | $\{AAD, ADD, CAD, CBB, CBD, CCB, CCD, CDD\}$ |
| 2 | $\{AB, BC\}$ | 13 | 6 | $\{AAC, AAD, BAC, BAD, BBD, CAC, CAD, CBD, CCD, DAC, DAD, DBD, DCD\}$ |
| | $\{AB, AC\}$ | 12 | 24 | $\{AAD, BAD, BBC, BBD, CAD, CBC, CBD, CCD, DAD, DBC, DBD, DCD\}$ |
| | $\{AB, CD\}$ | 12 | 1 | $\{AAC, AAD, ACC, ADC, ADD, BAC, BAD, BBC, BBD, BCC, BDC, BDD\}$ |

### D. Optimal doublet-triplet codes

Now we are ready to find the optimal doublet-triplet codes – the most promising doublet-triplet predecessors of the modern code.

First note that a comma-free doublet code (based on the 4-letter alphabet A,B,C,D) must have at least two sense doublets (*n*_2_ *≥* 2) to encode a non-trivial message, and can have at most five sense doublets [76] (*n*_2_ *≤* 5). Moreover, we can ignore the case *n*_2_ = 5, since from Table I it permits at most *n*^max^ = 2 compatible sense triplets, and it is unlikely that a code which can encode 5 AAs would be overtaken by a code which can only encode 2. Thus, we can narrow the cases of potential interest down to the cases where the initial doublet code had 2, 3 or 4 sense doublets.

#### 1. Assuming two-step evolution

First consider the optimal codes in the case of two-step evolution. In this case, the modern code evolved directly from some doublet-triplet code (with the triplet partner a subcode of the modern code). In this case, we find that the optimal doublet-triplet codes are the following:

- If the initial doublet code had four doublets, the five optimal doublet-triplet codes are shown in Fig. 5. The first three of these (red, orange, yellow) are the codes that, of those that maximize the number of distinct prebiotic AAs (6), also maximize the number of distinct Class-2 AAs (5), while the remaining two (blue, purple) are those that maximize the number of distinct Class-2 AAs (6) (of which 5 are then automatically Class 2).
- If the initial doublet code had three doublets, the six optimal doublet-triplet codes are shown in Fig. 6. The first four of these (red, orange, yellow, green) are the codes that maximize the number of distinct prebiotic AAs (8) (and they also then automatically code for a total of 9 distinct AAs, of which 6 are Class 2, and at least 5 are both prebiotic and Class 2), while the remaining two (blue, purple) are the codes that maximize the number of Class-2 AAs (7) (and they also then automatically code for a total of 9 distinct AAs, of which 7 are prebiotic, including all 6 AAs which are both prebiotic and Class 2).
- If the initial doublet code had two doublets, the five optimal doublet-triplet codes are shown in Fig. 7. The first of these (black) is the code that, of those that maximize the number of distinct prebiotic AAs (9), also maximizes the total number of distinct AAs (10), while the remaining four (red, orange, yellow, green) are the codes that, among those which maximize the number of Class-2 AAs (8), also maximize the total number of distinct AAs (12).

**FIG. 5.**
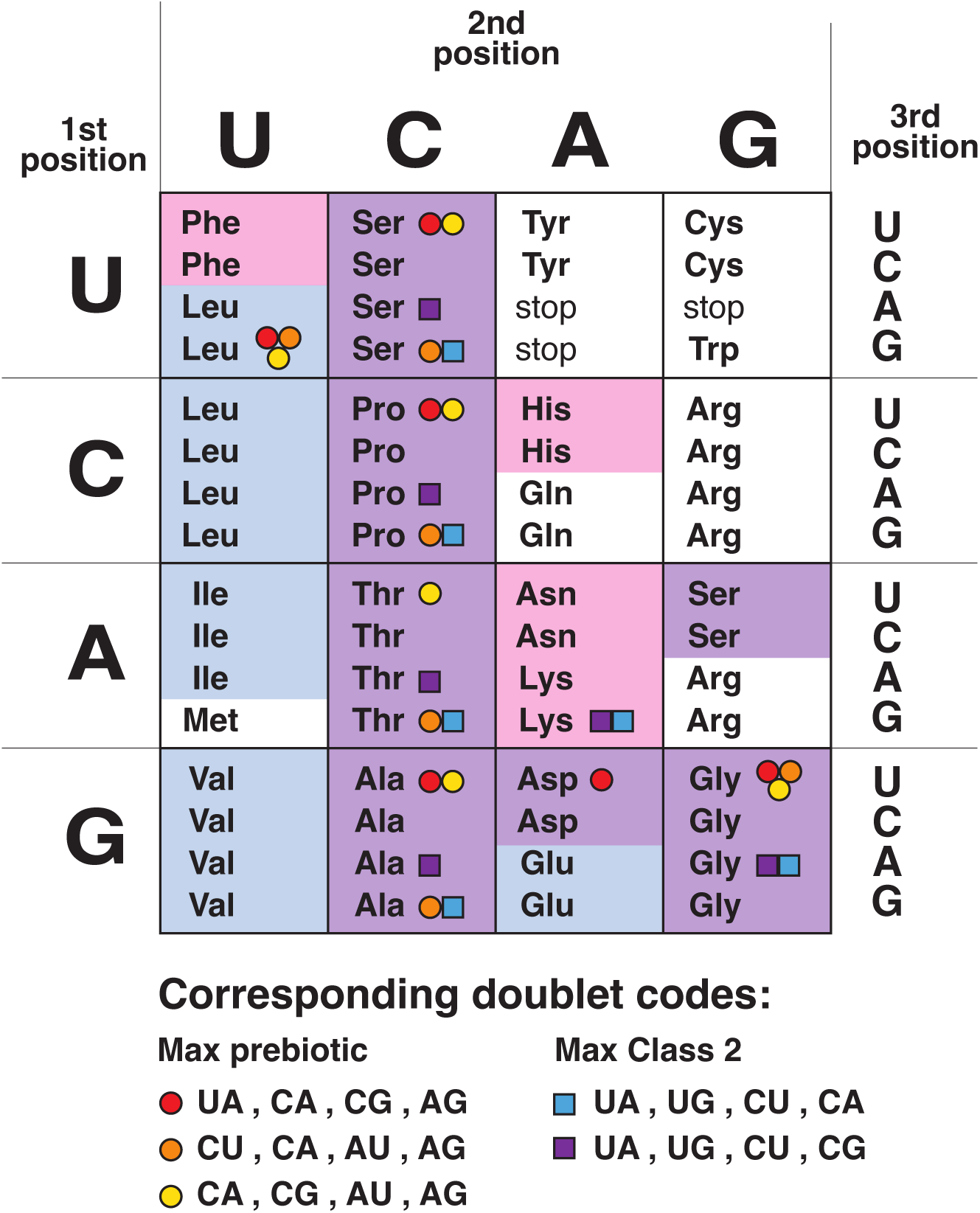
The optimal two-step doublet-triplet codes (assuming a four-doublet code).

**FIG. 6.**
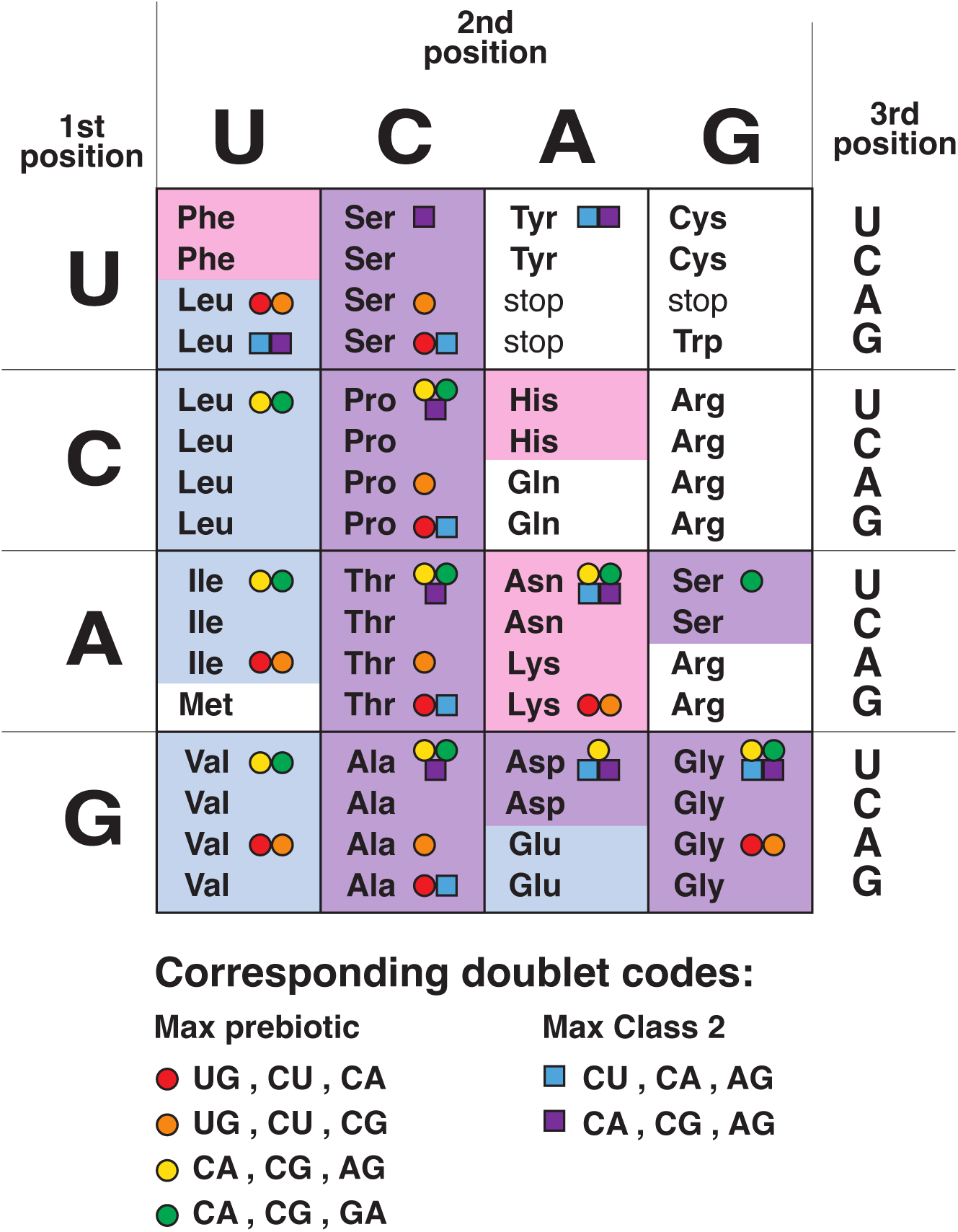
The optimal two-step doublet-triplet codes (assuming a three-doublet code).

**FIG. 7.**
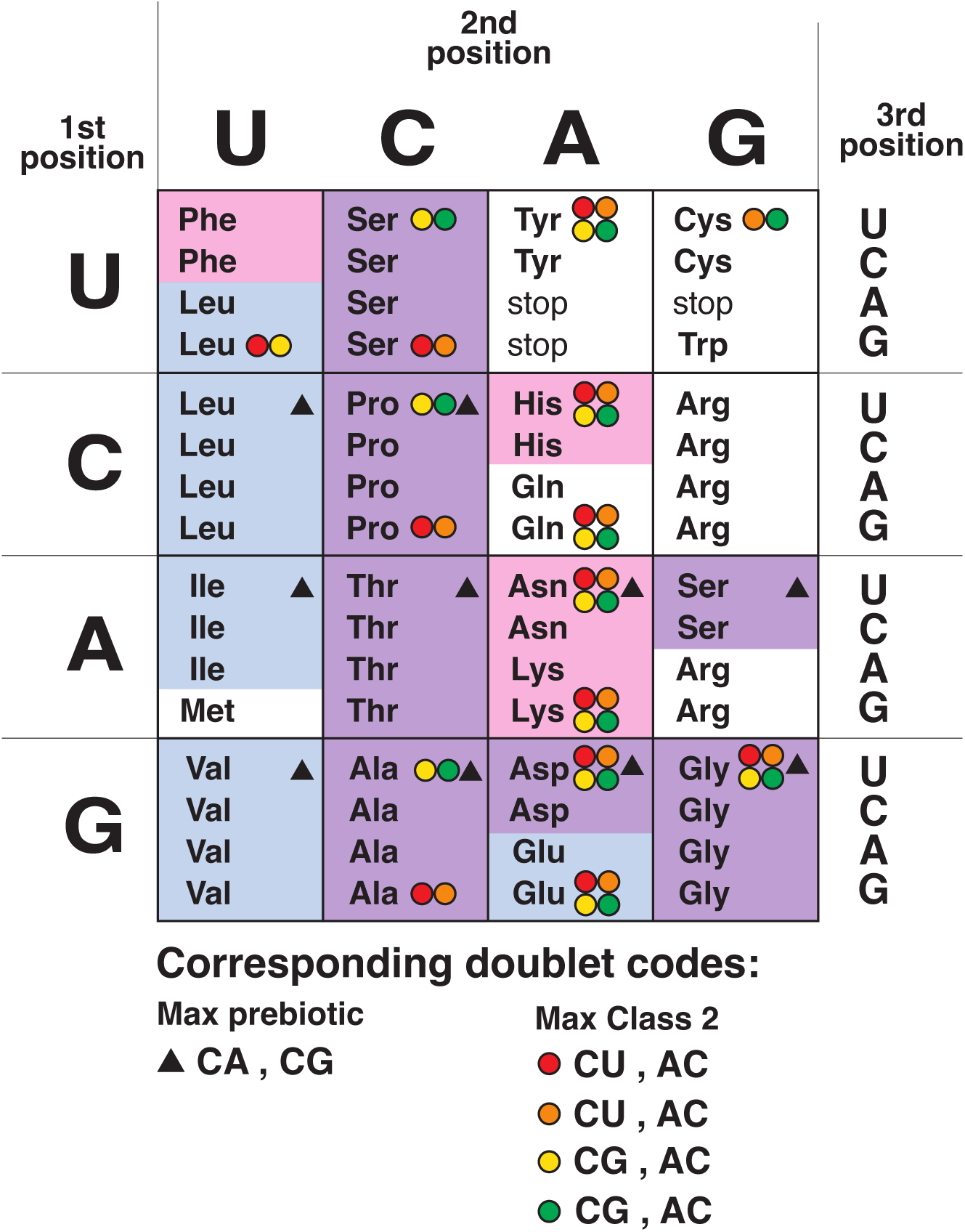
The optimal two-step doublet-triplet codes (assuming a two-doublet code).

#### 2. Assuming three-step evolution

Next consider the optimal codes in the case of three-step evolution. In this case, as explained above, the final comma-free triplet subcode grew from an initial doublet-triplet “seed code” whose doublet code became vestigial or disappeared. (For simplicity, we also focus here on the most interesting case where, in “Step 0”, the doublet code grew to the maximal size that still permitted it to subsequently be overtaken by a compatible comma-free triplet partner, *i.e.* a four-doublet code.) In this case, we find that there are three optimal comma-free subcodes:

- The first two optimal subcodes are shown in Fig. 8: they are very similar to one another, and each contain 14 distinct AAs. These are the two codes that, of those that maximize the number of distinct Class-2 AA’s (9), also maximize the total number of distinct AA’s (14) (and they then also automatically include 8 distinct prebiotic AAs, including all 6 AAs which are both prebiotic and Class 2). As indicated in the figure, these two codes grew, in turn, from one of five possible optimal doublet-triplet seed codes (which originally included a maximal number of distinct prebiotic or Class-2 AAs, before the doublet code became vestigial).
- The third optimal subcode is shown in Fig. 9: it is the code that, of those that maximize the number of distinct prebiotic AA’s (9), also maximizes the total number of distinct AA’s (12) (and it then also automatically include 8 distinct Class-2 AAs, including all six AAs which are both prebiotic and Class 2). As indicated in the figure, this code grew, in turn, from one of four possible optimal doublet-triplet seed codes (which originally included a maximal number of distinct prebiotic AAs, before the doublet code became vestigial); and, since it relates to a number of other interesting proposals about the the origin of the code (*e.g.* [14, 29]), we also note that it could have grown from a doublet-triplet seed code with just four triplets on the bottom row of the code (as case indicated by the blue triangles in the figure).

**FIG. 8.**
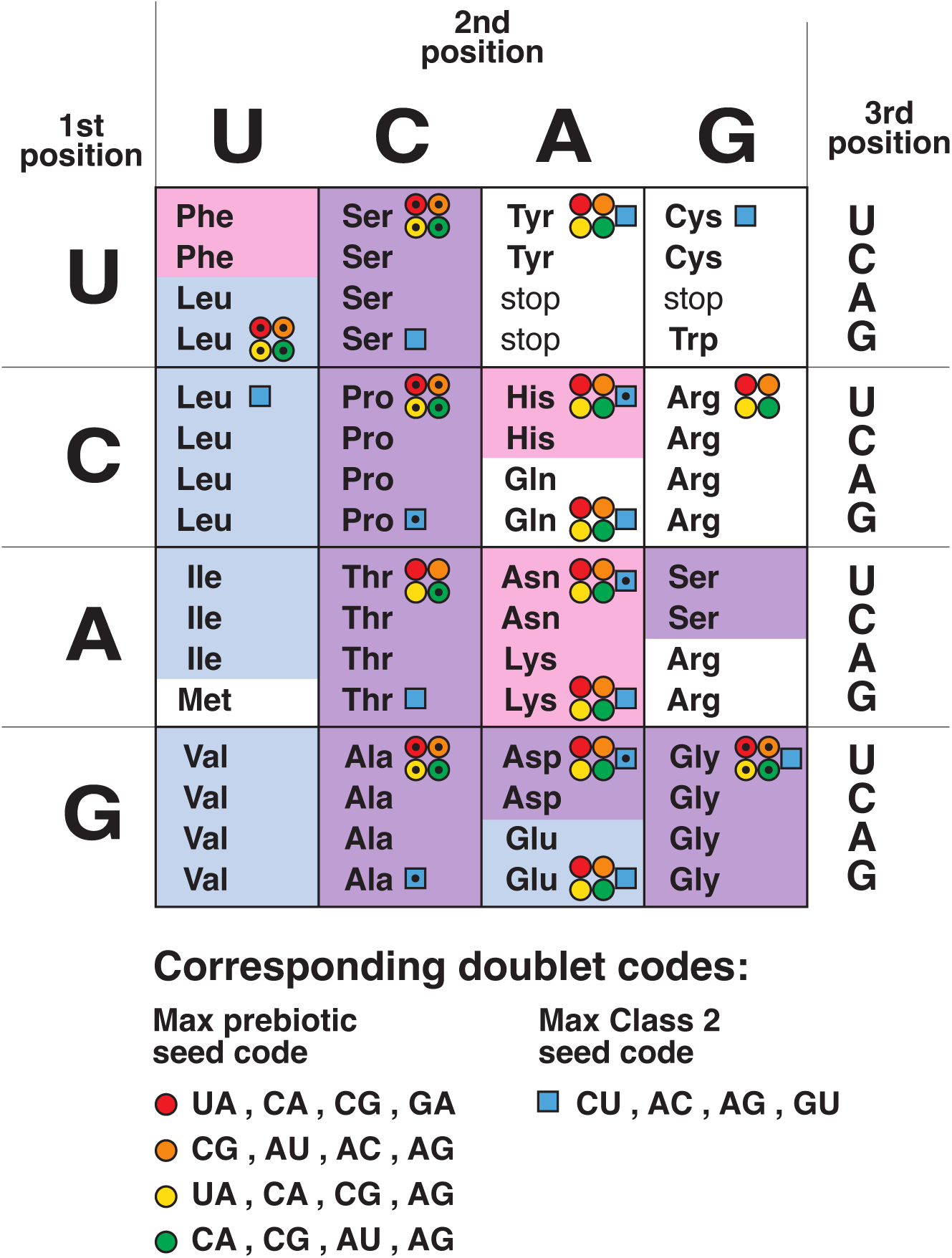
The two optimal three-step codes with 14-triplets, and the five optimal doublet-triplet “seed codes” from which they could have grown. (The seed-code triplets are indicated by interior black dots.)

**FIG. 9.**
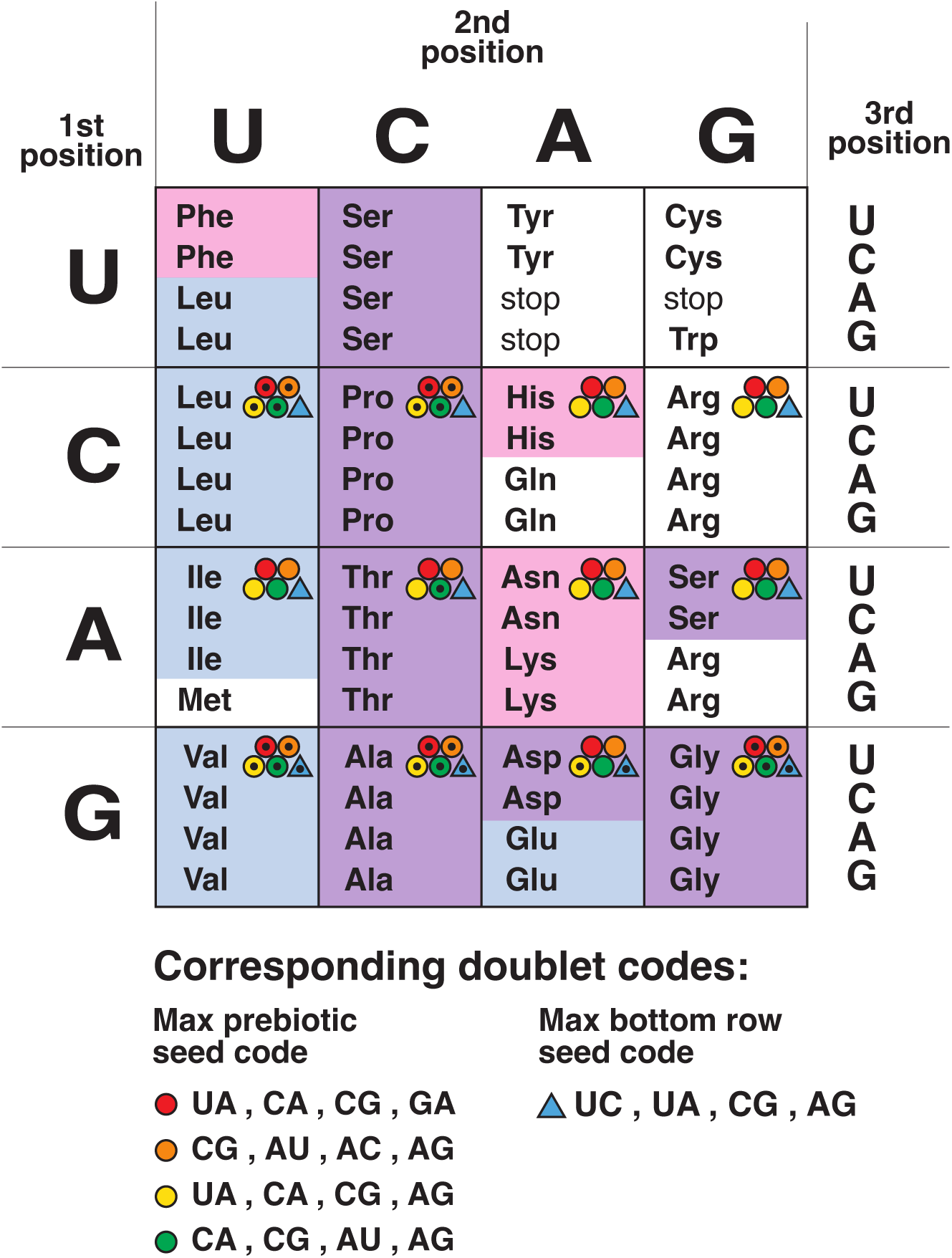
The optimal three-step code with 12-triplets, and the five optimal doublet-triplet “seed codes” from which it could have grown. (The seed-code triplets are indicated by interior black dots.)

## IV. DISCUSSION

### A. Including wobble, intriguing evidence

So far, all our analysis has implicitly assumed that the standard Watson-Crick pairing rules (A with U, and C with G) are strictly obeyed between the bases in the codon and the corresponding bases in the anti-codon. In reality (at least in the modern code, but possibly also in the primordial code) the pairing exhibits a phenomenon known as “wobble” [101–103]: in a triplet codon, although the first two codon bases follow strict Watson-Crick pairing with their corresponding anti-codon bases, the *third* codon base and its the corresponding anti-codon base (conventionally numbered as the first anti-codon base) follow a more flexible set of rules known as the wobble rules, summarized in Fig. 10.

**FIG. 10.**
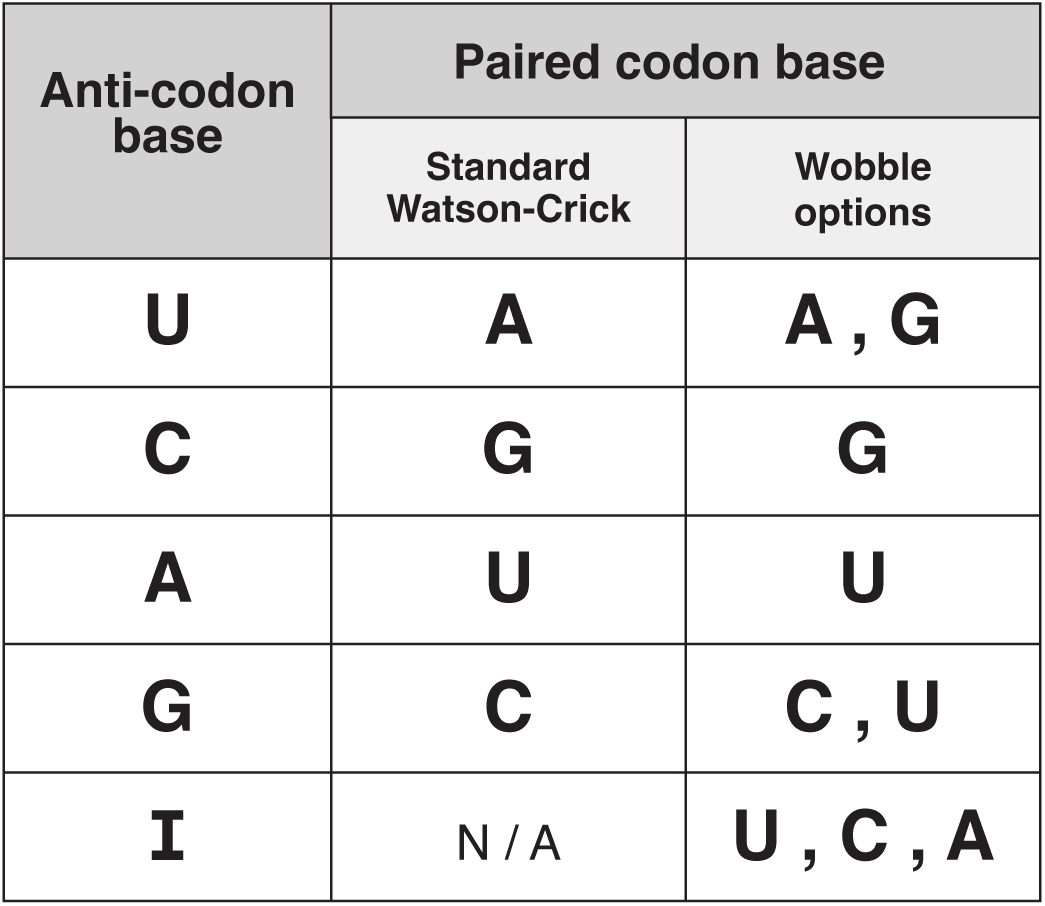
The standard wobble rules.

Let us understand how the notion of a comma-free code extends in the presence of wobble. In the absence of wobble, there is a one-to-one mapping between codon and anti-codon, so the two need not be specified independently. However, in the presence of wobble, we must be more careful: a given codon might be able to bind to more than one possible anti-codon, and vice versa.

Recalling that the underlying logic behind a comma-free code is to provide a *local* mechanism to unambiguously determine the correct reading frame, we see that in the presence of wobble, a comma-free code is one in which, for each sense codon, there *exists* a corresponding adaptor molecule (anti-codon) that can bind to it, and for any adjacent pair of sense codons, the adaptor cannot bind to the pair in the wrong reading frame.

To understanding how the problem of checking whether a given code (a given set of sense codons) is comma-free changes in the presence of wobble, consider Fig. 11. If the code includes a sense codon of the form XYU (where X and Y are arbitrary RNA bases) then, from Fig. 10, we see that there are three different anti-codons that could bind to it in principle (X YA, X YG, or X YI); but if the *actual* adaptor molecule that binds to it has anti-codon X YA, then this adaptor can *only* bind to the codon XYU. By contrast, if the code includes a sense codon of the form XYC, the adaptor molecule that binds to it (*e.g.* X YG) necessarily has an anti-codon which, in turn, can bind to multiple codons (in this case XYC or XYU).

**FIG. 11.**
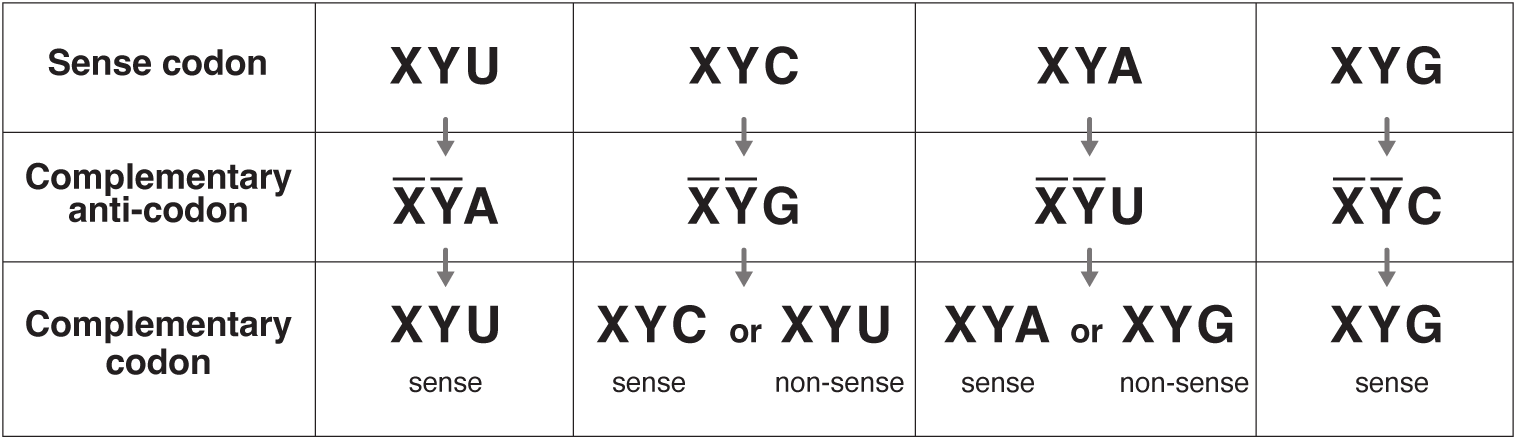
How to check a code is comma-free, including wobble effects.

Following this logic, one finds that nearly all codes that *would have been* comma-free, if one assumed strict Watson-Crick base pairing, are no longer compatible comma-free when the wobble is taken into account. In particular, if *all* the sense codons in a triplet code end in either U or G (denote such codons “Type 1”), then if the code is comma-free candidate when strict Watson-Crick pairing is assumed, it *remains* comma-free when wobble is taken into account. (We say such a code is “purely Type 1”.) But if *any* of the sense codons in a triplet code end in either C or A (denote such codons “Type 2”), then (for a comma-free code with sufficiently many sense codons) it will likely fail to be comma-free once the wobble rules are taken into account (especially for comma-free codes with many sense codons).

Thus it is striking that, despite the fact that the analysis in the previous sections assumed the strict Watson-Crick pairing rules (which carry no information about the wobble rules, and give no preference for Type 1 over Type 2 triplets), our three best candidates for the comma-free predecessor of the modern code (*i.e.* the three different three-step doublet-triplet codes identified in Sec. III D and displayed in Figs. 8, 9) all happen to be “purely Type 1” codes (*i.e.* all 14 codons in each of the first two codes, and all 12 codons in the third code happen to be Type 1 codons), so that these codes *automatically remain comma-free even when wobble is taken into account*!

As an (admittedly rough) argument to appreciate why this is striking, note that, on the one hand, a codon chosen at random has equal probability of being Type 1 (ending in *{*U,G*}*) or Type 2 (ending in *{*A,C*}*), so the probability that 14+12 triplets chosen at random would all be Type 1 is 2*^−^*^(14+12)^ *≈* 1.5 *×* 10*^−^*^8^.^3^ On the contrary, when we search for optimal comma-free subcodes hiding within the modern code, we appear to be led to subcodes that are purely Type 1 (with much higher probability than would be expected for a random subcode). Indeed, the preference for purely-Type-1 codes is pervasive throughout the various types of optimal subcodes found in Sections II and III: in particular, 4 of the 5 codes in Fig. 2 (orange, yellow, green, black), 3 of the 5 codes in Fig. 5 (red, orange, yellow), 4 of the 6 codes in Fig. 6 (yellow, green, blue, purple), all 5 of the codes in Fig. 7 (as well as the codes in Figs. 8, 9) are purely Type 1.

The fact that optimal comma-free subcodes of the modern code turn out to be Type 1 with much higher probability than would be expected for a randomly chosen subcode (so that these codes remain comma-free even when the effects of wobble are taken into account) appears to provide significant evidence for the possibility that the modern saturated code had a comma-free predecessor (and particularly significant evidence for the possibility of a three-step doublet-triplet predecessor). To this, we can add the various other supporting observations noticed previously (such as the fact that, in Sec. II C, the four comma-free codes that encoded the largest number of distinct amino acids (14 of 20) automatically also encoded the largest number of distinct prebiotic amino acids (9 of 10)). Systematically determining more accurate quantitative values for the strength of these various lines of evidence (separately and in conjunction with one another) is an important problem that we leave to follow-up work.

### B. A simpler starting point?

In Section III B, we explained how the idea of a compatible comma-free code could allow our triplet code to have been preceded by a simpler doublet code. It is interesting to consider the possibility of other, even simpler, starting points.

The simplest starting point would be a singlet code (where each codon is a single letter), but this is not a promising possibility, since a singlet code and a more complex (doublet or triplet) code can never be mutually compatible (and hence do not satisfy the local constraints to allow them to evolve in parallel without interfering with one another) unless the two codes use two completely distinct, non-overlapping alphabets.^4^

The next simplest starting point would be a doublet code that only uses one of the two RNA base pairs (either *{*A,U*}* or *{*C,G*}*, but not both); but again this doesn’t work, because a comma-free doublet code based on a two-letter alphabet can only contain a single sense doublet, so cannot encode a non-trivial message.

Interestingly, by this line of reasoning, the starting point analyzed in Sec. III (a comma-free doublet code based on two pairs of letters: *{*A,U*}* and *{*C,G*}*) is the simplest starting point that works!

It is, however, worth mentioning another, comparably simple starting point: namely, a triplet code based on just two letters (*i.e.* one base pair, *e.g. {*C,G*}* (or *{*A,U*}* or even *{*A,I*}*, as originally suggested by Orgel [9]). A comma-free triplet code based on two letters can include at most two sense codons which, from a logical standpoint, is enough to encode a non-trivial message, although (with a repertoire of only two different amino acids) it is unclear whether it has enough chemical flexibility to produce useful proteins. In any case, if life could have started with such a code, it would indeed permit a nontrivial compatible comma-free triplet code based on four letters. So the path from this starting point to the current triplet code does seem to be a logical possibility in principle, although we have not yet found any notable arguments or evidence in its favor.

### C. Final remarks

In this paper, we explained a general reason to expect that the primordial genetic code was comma free, studied the possibility that the modern saturated triplet code evolved from an earlier comma-free triplet code, and identified the five optimal candidates for such a primordial triplet code. We then introduced the idea of a “compatible comma-free code,” explained how it could have allowed a primordial comma-free triplet code to have evolved from an earlier comma-free doublet code and again, for each of the various possible evolutionary paths, identified the optimal candidates for these primordial doublet and triplet codes. Finally, we explained how the so-called wobble rules affect the story, and how the striking compatibility of the optimal codes with these wobble rules seems to provide non-trivial evidence for this scenario.

We end by pointing out that each of our optimal codes makes detailed predictions about the earliest codons in the code, and the order in which they entered. By comparing these predictions to other lines of evidence about the order in which various codons and AAs might have entered the code (*e.g.* [11, 26, 47, 55–67, 90, 92, 93, 104–107]) it may be possible to make further progress in sorting through these possibilities and thereby obtain more detailed clues about the pathway that the early evolution of the code followed. This is a very interesting direction for future work.

## Supporting information

Mathematica code for generating doublet-triplet codes

Mathematica code for comparing comma-free subcodes to modern code

## Acknowledgements

We thank Elina Mer for careful reading and help with the figures. Latham Boyle is supported by the STFC Consolidated Grant ‘Particle Physics at the Higgs Centre’. Research at Perimeter Institute is supported by the Government of Canada, through Innovation, Science & Economic Development, Canada and the Province of Ontario through the Ministry of Research, Innovation & Science.

## AI usage statement

AI was used for finding some relevant citations, but was not otherwise used in the research or writing of this paper.

## Footnotes

1 Henceforth in this paper, when we refer a gene and its constituent codons, we will always be referring to the RNA version of the gene. In the modern code, this is an mRNA copy of the primary DNA version; and at earlier times (if the RNA-world hypothesis is correct [9, 82–87]) the RNA version may have been the primary one.

2 That is, if *c^′^*was previously a sense codon, the fact that *c^′^*is now translated by two competing adaptors (*c^′^, x*) and (*c^′^, x^′^*) may introduce additional mutations.

3 One may worry that this argument is *too* rough, since it effectively treats the 14+12 codons as independent, whereas they are in fact correlated through the comma-free restriction. This is certainly true, but as one can see from the abstract classification of maximal comma-free codes in [77], the comma-free condition on its own does *not* cause the codons to all share one or two final bases; rather, even maximal comma-free codes typically include all four final bases.

4 Another more speculative problem with a singlet-code starting point is that, just as one would naively expect a doublet codon and its anti-codon to bind less reliably than a triplet codon and its anti-codon (because the triplet case has three RNA base-pairs bonding, while the doublet case has only two), one would expect a singlet codon and its anti-codon to bind even less reliably (since it has only one base-pair bonding).

